## Supplemental for "RecombiCraft Library construction: A novel method for DNA Library cloning and expansion using non-enzymatic single-step DNA recombination and liquid culture"

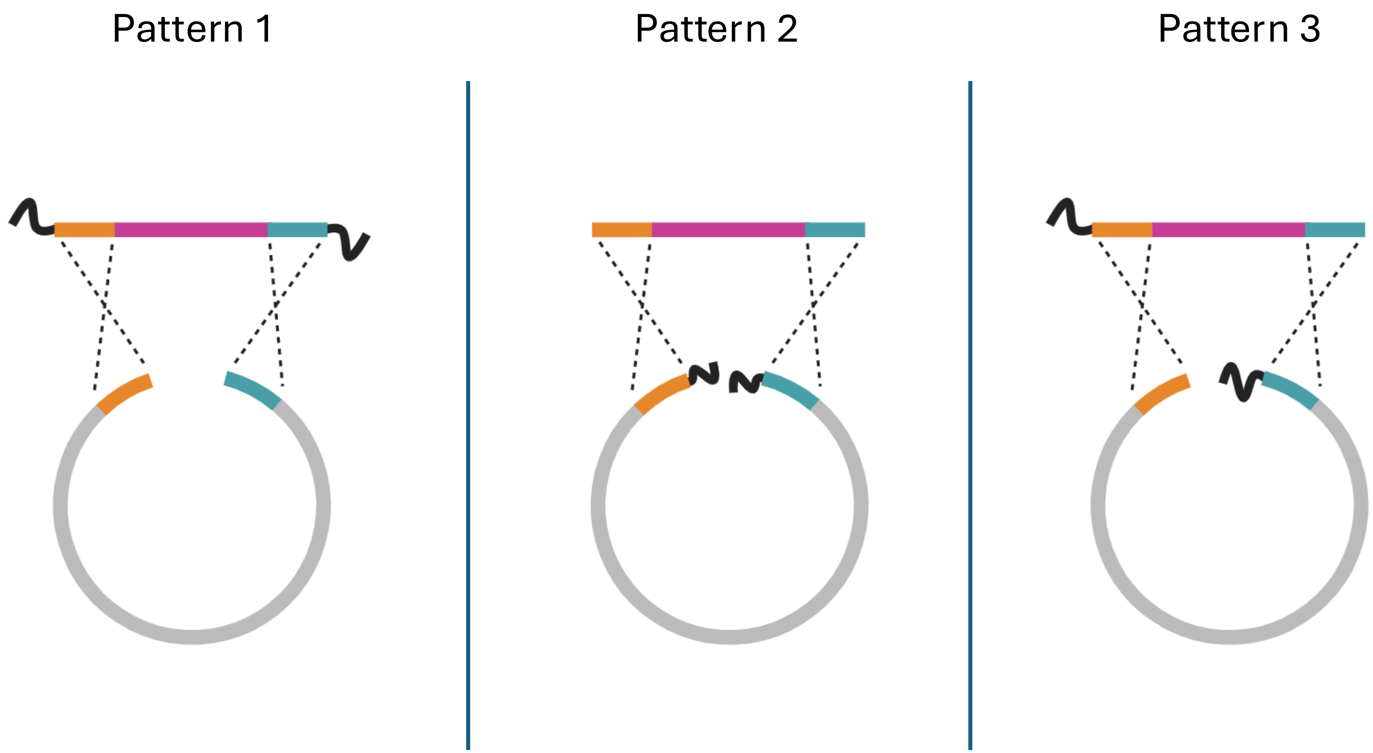


**Supplemental Fig S1.** The design flexibility of SLiCE cloning. SLiCE can flexibly assemble the insert and backbone with flanking sequences.

Supplemental Table S1

| Primer | Sequence (5’-3’) |
|---|---|
| Lib-insert-F | CCAGCCTCCTCGTCGCCTAT |
| Lib-insert-R | CGATCCGCCACCGCCAGA |
| Lib-backbone-F | GGTGGAGGCGGTTCAGGC |
| Lib-backbone-R | AGCATAATCTGGAACATCATATGGATAGGC |
| Seq-F | ACACTGACGACATGGTTCTACACAACGCAACTCCGAACTCT |
| Seq-R | TACGGTAGCAGAGACTTGGTCTGGTGCGATAATTGATGCTGATTG |

**Supplemental Table S1. The list of primers used in this study**
